# MeTrEx: Membrane Trajectory Explorer

**DOI:** 10.1101/2025.01.31.634501

**Authors:** Sabrina Jaeger-Honz, Christiane Rohse, Beat Ehrmann, Karsten Klein, Ying Zhang, Wendong Ma, Yu Hamada, Jian Li, Falk Schreiber

**Affiliations:** Department of Computer and Information Science, University of Konstanz, Universitätsstrasse 10, Konstanz, 78464, Germany; Monash Biomedicine Discovery Institute, Monash University, Clayton, 3800, Victoria, Australia; Faculty of Information Technology, Monash University, Clayton, 3800, Victoria, Australia

**Keywords:** Molecular Dynamics Simulation, Membrane Interaction Analysis, Trajectory Visualisation, Ligand-Membrane Interaction, Visual Analysis, Computational Biophysics

## Abstract

**Background:** Molecular Dynamics (MD) simulations provide valuable insights into the behaviour of biomolecules, particularly in complex systems such as cellular membranes. However, the increasing volume of MD data presents significant challenges in data visualisation, exploration and analysis.

**Result:** We present MeTrEx (Membrane Trajectory Explorer), a Python-based software designed to enhance the exploration and analysis of MD simulation data, with a focus on ligand and membrane molecule interactions. MeTrEx offers an intuitive graphical user interface for visualisation, advanced interaction tools, and comprehensive analytical capabilities. The software also supports comparative analyses of molecular trajectories and integrates external datasets for a more holistic view. A case study analysing the interaction of Polymyxin B1 with a bacterial membrane demonstrates the tool’s capabilities to investigate complex molecular behaviours.

**Conclusion:** By combining detailed molecular analysis with interactive visualisation, MeTrEx provides researchers with a powerful platform to investigate complex biomolecular systems, contributing to a deeper understanding of ligand-membrane interactions.

## 1 Background

Molecular Dynamics (MD) simulations aid in understanding and modelling biomolecules such as cell membranes to understand their 3D conformations, interactions with other biomolecules, or penetration [1, 2]. MD simulations result in massive time series data sets with individual atoms and their coordinates at a particular time, so-called trajectories [3]. While more and more data can be routinely calculated and collected, a significant bottleneck for MD simulations is the subsequent data analysis. Visualising and analysing long trajectories becomes increasingly difficult as the size of the trajectories increases due to the computational requirements and the need to create interactive representations that facilitate human reasoning within the limits of human perception and cognition [4–6]. Good analysis software should identify potential points of interest and guide the analyst to them, avoiding the need to look up all calculated data points.

In a typical analysis workflow, points of interest are identified by calculating established measures, such as root-mean-square deviation, root-mean-square fluctuation, radius of gyration or energy-based approaches [7, 8]. These points of interest can then be visually inspected and explored.

In recent years, MD simulations have become more powerful and accessible with improved speed and accuracy. Accordingly, a large number of tools have been developed to analyse MD simulation data [9]. Some tools focus on visualisation and interaction but also provide basic analysis methods (e.g., Visual Molecular Dynamics [10], PyMOL [11], ChimeraX [12]), others offer many possibilities for general calculations or interaction (e.g., Travis [13], MDTraj [14], MDAnalysis [15], mdciao [16]). Various specialised tools emerged to provide domain-specific calculations and focused on specific aspects of MD simulation. One group of tools specialises in membrane MD simulations because of the increased interest in studying membranes. Some of them focus on protein-membrane simulations (e.g., PyLipID [17], Prolint [4], DipTool [18]), others focus solely on the membrane (e.g., LiPyphilic [19], APL@voro [1], Membrainy [20], MemSurfer [21], MOSAIC [22]).

While current tools for membrane MD analysis offer valuable computations, they often lack advanced membrane-molecule interaction capabilities. For instance, ProLint and APL@voro provide fundamental analysis but with limited user interaction features [1, 4]. Tools like MemSurfer and Membrainy focus on specialised analyses, but their command-line interfaces and non-interactive visualisations limit accessibility [20, 21]. Similarly, DipTool, designed for quantitative structure-activity relationship (QSAR) based ligand-membrane interaction studies, is incompatible with MD simulation data and lacks comprehensive visualisation features [18]. Overall, these tools are unsuitable for detailed ligand-membrane interaction studies and do not offer the necessary interactivity and visual exploration for complex datasets. Table 1 compares the most commonly used and recently published tools and their features.

**Table 1:**
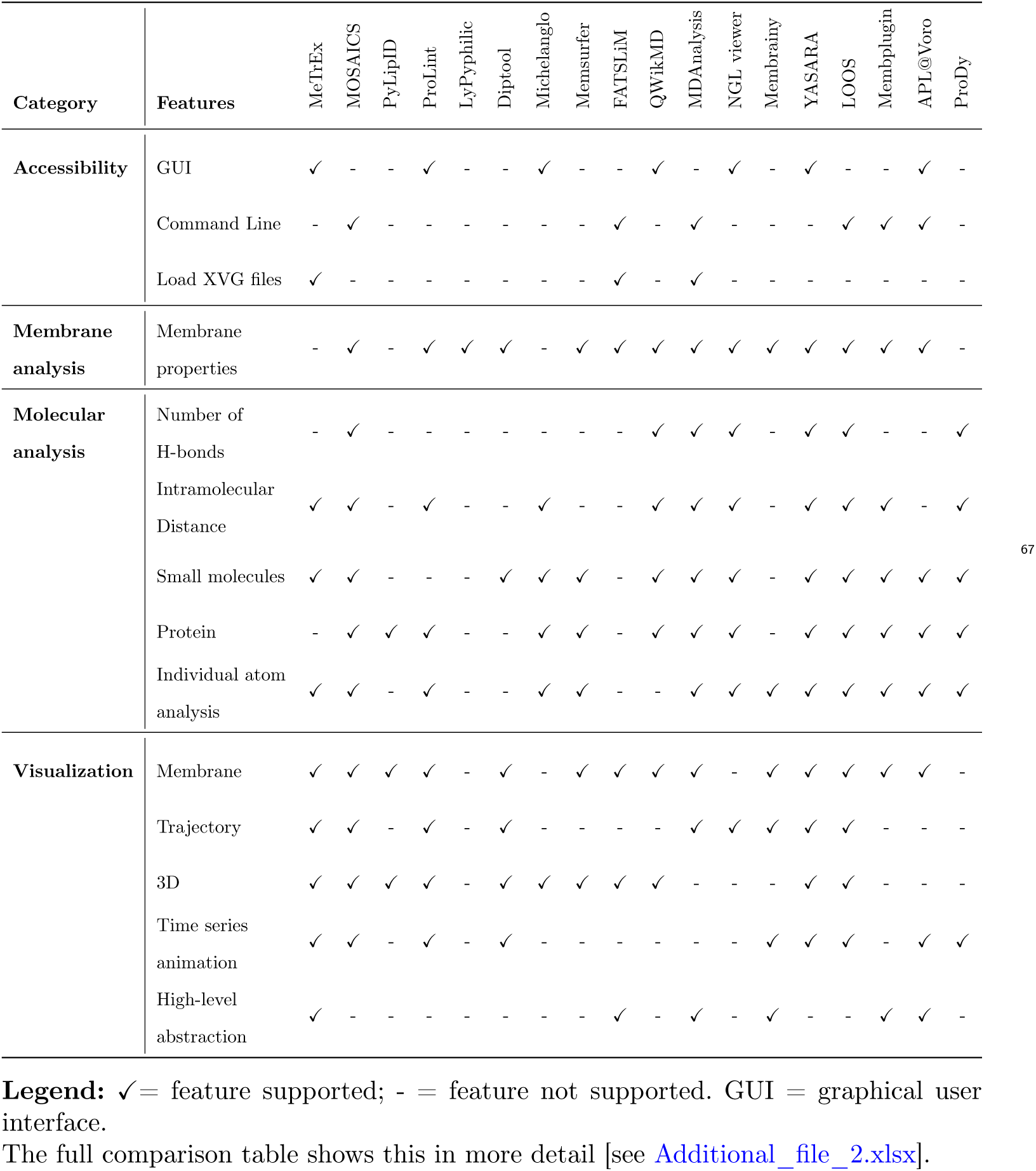
Summary of tools and their features for analysing and visualising MD simulation data.

To address the growing complexity of MD simulation data, researchers require tools that go beyond basic visualisation and offer advanced interaction, comparison, and sharing capabilities. Programs like sMolBoxes provide valuable features such as data provenance visualisation and guided user interaction, but they are tailored for protein-based studies and lack membrane-specific analyses [23]. MOSAIC is written in C++ and designed for advanced analysis of lipid layer structure [22]. LOOS offers a higher performance by combining C++ and a Python library designed to enable rapid development for MD analysis. However, these two have some limitations: For example, most analyses are provided as 2D distributions and require the integration of additional software like VMD or PyMOL to get 3D visualisation. There is a clear need for an integrated tool combining comprehensive membrane and ligand interaction analyses with advanced visualisation features in a user-friendly, interactive environment. Developing such a solution would significantly enhance the exploration and understanding of complex biomolecular systems [18].

We designed MeTrEx (Membrane Trajectory Explorer) to enable the visual exploration and analysis of MD simulation data, particularly emphasising membrane and ligand interactions. Its primary aims include providing researchers with intuitive, interactive visualisation for exploring complex MD datasets and enabling real-time manipulation and investigation of molecular trajectories. MeTrEx is intended to support in-depth analysis by offering a range of analytical techniques, including the calculation and visualisation of intramolecular distances, molecular speeds, and other key properties. Additionally, the software promotes comprehensive data integration by seamlessly incorporating diverse data formats and external analysis files into the workflow. To ensure accessibility for a wide range of users, MeTrEx offers a user-friendly interface, enabling even those without extensive technical expertise in MD to efficiently perform complex analyses.

## 2 Implementation

MeTrEx is optimised to combine an overview visualisation and a set of analytic tools for MD simulations aimed at ligand-membrane systems in an easily accessible graphical user interface (GUI) (see Fig. 1). To import simulation data, MeTrEx uses a range of file formats for structure data (i.e., Worldwide Protein Data Bank format PDB and GROMACS format GRO) and trajectories (i.e., GROMACS compressed trajectory format XTC and GROMACS TRR trajectory format). The XVG format, the default output file format of the GROMACS molecular dynamics package, is also supported for visualisations of additional external analyses [24]. After the 3D overview visualisation is initialised, MeTrEx offers a set of tools to map frame positions, molecular speeds, and intramolecular distances onto molecular trajectories. In the following, we will denote the central view of our system with the main visualisation of the trajectories and membrane as the main view and subwindows derived from the main view as sub-views. Interactive exploration tools of the main view, including panning, zoom controls, time-frame navigation sliders, and the ability to isolate specific trajectories, empower users to investigate regions of interest in detail. For more granular analysis, MeTrEx provides the so-called bottom view displays. In these, detailed calculations of atomic distances and molecular speeds are shown in additional graphs below the main view, supporting both single-atom and multiple-atom/molecule analyses. These features provide precise control over the visualisation, enabling detailed analysis of spatial relationships, molecular interactions, and temporal changes within the membrane environment. The software also facilitates the integration of external data through the sub-views, enabling users to visualise supplementary datasets alongside their simulation results. Additionally, for data provenance visualisation, every molecule trajectory in the main view is assigned to a single colour, which is used as a preset for the corresponding analysis visualisations in the bottom views and, if selected by the user, in the sub-views. All supported colour gradients are perceptually uniform, and schemes for colour-vision-deficient and colour-blind people are included. Finally, users can save and export their findings, including atom positions, frame selections, and visualisations, for further use or collaboration.

**Figure 1:**
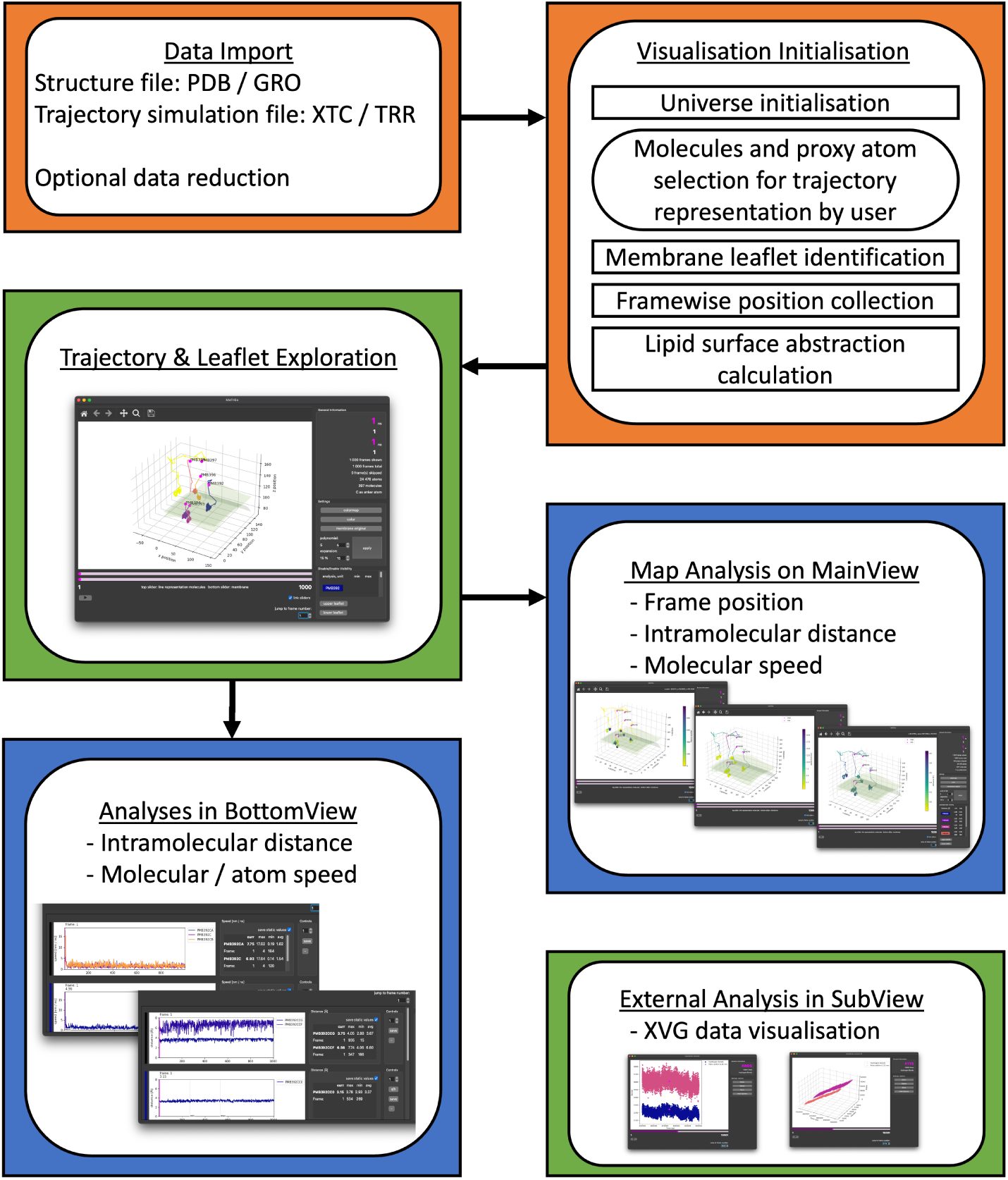
Flow chart of the functionality of MeTrEx. **Initialisation (orange)**: To start the visualisation, both structure and trajectory-simulation data must be provided. **Visualisation (green)**: The 3D visualisation in the main view of MeTrEx allows exploration of the molecular trajectories and membrane simulation. Supplementary XVG analysis data can be displayed in a separate instance called the sub-view. **Analysis (blue)**: In-depth analysis tools include mapping analysis onto the trajectories and graphical analysis visualisation within the so-called bottom view.

We developed MeTrEx as a Python-based standalone software to provide a robust and flexible platform for visual exploration and analysis of MD data, with system requirements including Python 3.8 or higher supported by Windows, Linux and Ma-cOS operating systems. The graphical user interface (GUI) is built using the *PyQt6* framework [25], incorporating *QtCore*, *QtGui*, and *QtWidgets* libraries for an intuitive, interactive experience, with a menu bar, toolbar, and well-structured layouts. For data handling and analysis, MeTrEx integrates the *MDAnalysis* [15] and *MDTraj* [14] libraries, widely used in the MD community to process and analyse trajectory data efficiently. Visualisation is powered by *Matplotlib* [26], enabling dynamic and customisable 2D and 3D visualisations of molecular trajectories. Lipid surface regression is specifically implemented using using the approach by Kern et al. [1], while *scikit-learn* is used to calculate surface abstractions [27]. MeTrEx employs a modular architecture based on the Model-View-Controller (MVC) design pattern [28], ensuring a clear separation of logic, visualisation, and data storage; this makes the application easier to maintain and extend. Performance is optimised through deferred updates, refreshing views only after user interactions to reduce computational overhead. Dialogues, message boxes, and clear visual feedback enhance user interactions. Robust error handling and informative messages guide users through potential issues.

## 3 Results

To illustrate the capabilities of MeTrEx in analysing MD simulations of membrane interactions, it was applied to a dataset simulating the behaviour of Polymyxin B1 (PMB) on a bacterial membrane. This analysis serves as a case study to demonstrate the utility of MeTrEx in visualising and interpreting ligand-membrane MD data, particularly for investigating ligand interactions (e.g., drugs such as PMB) with complex membrane systems. The dataset used here was initially published by Jaeger-Honz et al. to test MeTrEx, offering a foundation to study the molecular dynamics of PMB with a focus on its interactions with the membrane of *Pseudomonas aeruginosa*[29].

Antibiotic resistance is a critical global health issue, especially in bacteria like *Pseudomonas aeruginosa*. This pathogen is notorious for its resistance to multiple drugs and high transmissibility, posing a significant mortality risk worldwide, according to the World Health Organization [30]. Polymyxins, including PMB, are among the last-line antibiotics effective against *P. aeruginosa*. These antibiotics, distinguished by a peptide ring and an extended fatty acid or acyl chain, are often used when other antibiotics have failed to control infection [31–33]. Studying the interactions of polymyxin antibiotics with bacterial membranes at the molecular level is thus essential for understanding their mechanisms of action and potential optimisation strategies to counteract bacterial resistance.

### 3.1 Visualisation of Molecular Trajectories and Interaction with Membrane Surfaces

The main view of MeTrEx displays the trajectories of individual PMB molecules, with each molecule labelled by its residue number. This visualisation offers an initial high-level overview of molecular movements, allowing users to assess spatial patterns and track the trajectories of each molecule over time. This feature is invaluable for observing molecules’ general behaviour and distribution within the membrane environment. With the interactive elements of MeTrEx, users can focus on specific frames or animate the trajectory in a time-lapse sequence, providing insights into the temporal dynamics of molecule-membrane interactions (see Fig. 2A and Additional_file_1: Fig. S1).

**Figure 2:**
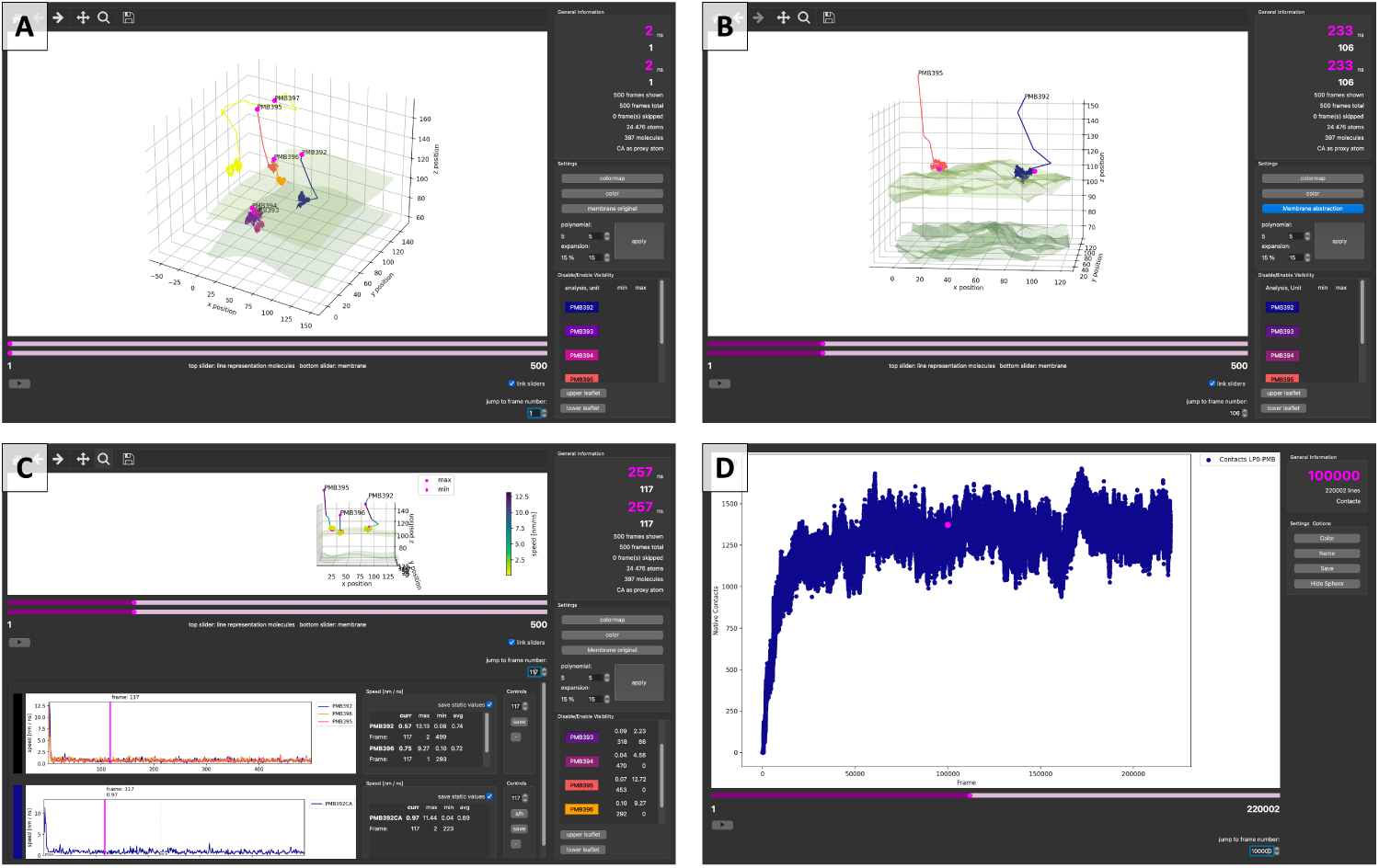
Comprehensive visualisation and analysis in MeTrEx: **A** The main view displays Polymyxin B1 (PMB) trajectories alongside the bacterial double lipid membrane. **B** Highlighting specific trajectories and temporal positions with interactive 3D tools (rotation, zoom, and pan). **C** Molecular speed mapped along trajectories, with focused analysis in the bottom view. **D** Sub-views visualise external XVG data in customisable 2D/3D plots in a subwindow.

Beyond basic trajectory visualisation, MeTrEx allows users to overlay molecular trajectories onto the membrane’s original structure, offering a realistic depiction of molecular proximity to various membrane regions. This less abstracted membrane visualisation, which uses the phosphor atoms of the lipid molecules as surface representation, is essential for directly observing molecular interactions at specific membrane sites. In addition, MeTrEx provides users with flexible viewing options such as rotation and zoom, enabling detailed observation of molecule orientation and interactions with the membrane surface from multiple angles. To further refine the visualisation, MeTrEx supports adjusting the polynomial degree of membrane abstractions, enhancing the level of detail and highlighting subtle interaction patterns between molecules and the membrane (see Fig. 2B and Additional_file_1: Fig. S2).

### 3.2 Mapping Molecular Positions, Distance and Speed Over Time

One of MeTrEx’s features is mapping molecular positions over time using a colour gradient onto the trajectories in the main view. This feature effectively visualises the progression of molecular movement, with the gradient indicating specific frames and allowing researchers to assess temporal distribution along the trajectory more easily (see Additional_file_1: Fig. S3).

Furthermore, the software can represent molecular or atomic speed through a colour gradient, quantifying the speed in nanometres per nanosecond (nm/ns). For instance, minimum and maximum speeds for each molecule are displayed, aiding in identifying periods of high molecular interaction with the membrane surface, which is essential in studying antibiotic-membrane interactions(see Fig. 3 and Additional_file_1: Fig. S4).

**Figure 3:**
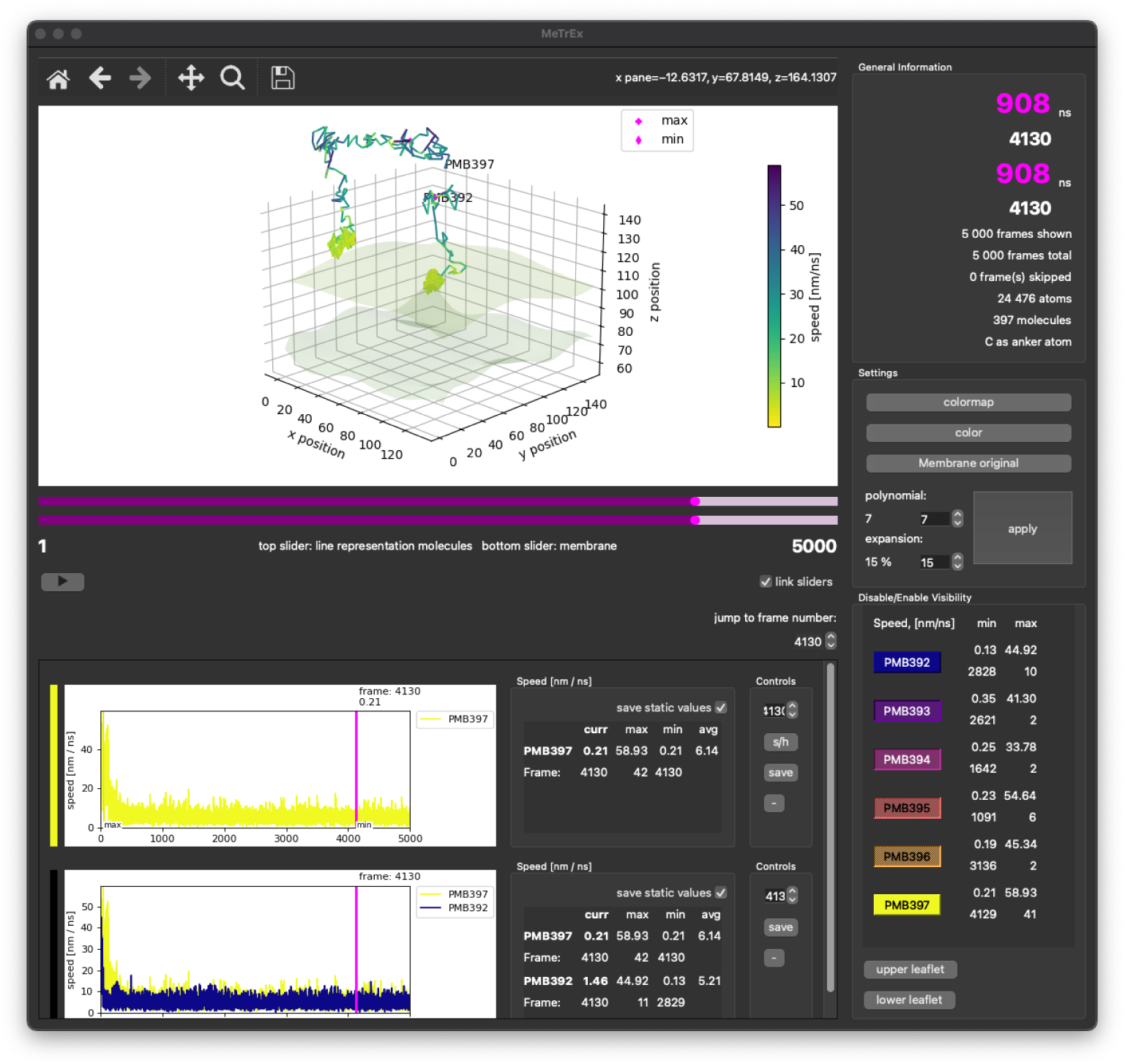
Comparison of molecular speed over time in the main view and bottom view: Molecular or atomic speed is visualised in the main view using a colour gradient mapped along the trajectories. In the bottom view, the upper graph depicts the speed of a single trajectory, while the lower graph compares the speeds of both displayed trajectories. Both views highlight that the molecules initially exhibit high speed, which decreases as they approach the upper membrane leaflet. Subsequently, the molecules anchor themselves into the membrane, resembling an insertion process.

In addition, MeTrEx can map molecular and atomic distances over time to the trajectories, providing an in-depth view of how molecules interact within each other or specific membrane regions. By tracking distance changes between molecules and the membrane, researchers can observe critical interaction dynamics, such as binding, penetration, and proximity to different membrane sites, which are essential for understanding ligand-membrane interactions. This distance-based visualisation enables precise identification of when and where a molecule comes into close contact with the membrane, shedding light on critical events like initial binding or deeper insertion into the lipid bilayer.

These capabilities are beneficial for observing how PMB molecules behave over time as they interact with the membrane and exert their effect. It potentially identifies moments of penetration or strong binding, which can be further investigated for drug efficacy analysis [34]. The visualisation of the example data illustrates how the simulated PMB molecules quickly interact with the upper leaflet of the membrane at the beginning of the simulation and subsequently anchor themselves within it (see Fig. 3).

Mapping atomic-level intramolecular distances over time to the trajectories allows for observing subtle interactions that may not be evident from broader trajectory visualisations. For instance, this feature can highlight sites of molecular bonding or *Van der Waals* interactions, which play a significant role in the behaviour of ligands like PMB within a bacterial membrane [35].

### 3.3 Focused and comparative analysis of molecular trajectories

MeTrEx provides powerful tools for both focused and comparative analysis of molecular trajectories, allowing users to gain detailed insights into individual molecule behaviours and interactions across multiple trajectories. For a more in-depth analysis, users can isolate the trajectory of a single molecule, such as PMB, and examine its speed over time within the bottom view. This detailed view plots molecular speed, displaying essential metrics such as speed at a selected frame, average speed, and minimum and maximum values, offering valuable data points for understanding individual molecular interactions with specific membrane regions. Especially when studying antibiotics like PMB, it is beneficial to see where interactions with specific membrane regions may contribute to their effectiveness (see Fig. 2C, Fig. 3 and Additional_file_1: Fig. S5).

MeTrEx enables the simultaneous display and comparison of multiple molecular trajectories for comparative analyses. In this mode, a bottom view shows combined graphs of molecular speeds across different molecules, allowing users to observe and compare speed and interaction frequency variations within the membrane environment. This comparative capability is advantageous for studying diverse interactions and identifying patterns across different molecules. The workspace remains userfriendly and organised even when multiple graphs are displayed, with scrollable options and the ability to remove graphs individually. This structured setup maintains efficiency, making MeTrEx a versatile tool for focused and comparative molecular trajectory analyses.

### 3.4 Visualisation of external data in sub-views

MeTrEx also supports external data visualisation through sub-views, which can be configured to display data from external XVG files. These sub-views enable users to import, customise, and analyse additional datasets in 2D or 3D scatter plots. Fig. 2D (and Additional_file_1: Fig. S6) show an example of this functionality, where external analysis data from an XVG file is displayed. Researchers comprehensively understand molecular dynamics by linking the imported data with the primary visualisation in the main view and the analytic findings in the bottom view. Sub-views further allow for interactive customisation, such as adjusting colours and renaming data labels, enhancing the accessibility and interpretability of imported data within MeTrEx.

## 4 Discussion and Conclusion

MeTrEx provides a comprehensive and flexible framework for visualising and analysing membrane MD simulations. It offers tools that are particularly well-suited for studying the molecular behaviours of ligands, such as antibiotics like PMB. The object-oriented architecture and Python framework allow new analyses or functions to be added easily. MeTrEx excels at linking visualisations and analyses, allowing researchers to view multiple properties and trajectories simultaneously in a unified framework. Integrating these aspects facilitates a more comprehensive understanding of MD simulation data. For instance, the visualisation and analysis of the example data illustrate how MeTrEx helps uncover key insights, such as how PMB molecules move towards and anchor in a membrane. Furthermore, researchers can explore trajectories alongside property analyses to identify interesting time points or patterns, whereas traditionally, they would need to check graphs separately and manually map key moments. This unified approach simplifies and accelerates the complex analysis of ligand-membrane interactions.

However, high rendering times when handling large datasets suggest that alternative visualisation libraries or GPU support could significantly improve performance. Future enhancements could include membrane-specific analyses, additional molecular analyses such as hydrogen-bond analysis, and 3D ball-and-stick visualisations for selected molecules.

To conclude, by enabling researchers to explore MD simulation data through interconnected visualisations and analyses, MeTrEx supports a more detailed and efficient investigation of ligand behaviours. These strengths establish MeTrEx as a valuable tool for the complex and comprehensive analysis of MD simulations, potentially improving understanding and driving discoveries in the field.

## Availability and requirements

- Project name: MeTrEx
- Project home page: https://github.com/LSI-UniKonstanz/MeTrEx
- Operating system(s): Platform independent
- Programming language: Python
- Other requirements: Python 3.8 or higher
- License: GPL3
- Any restrictions to use by non-academics: None

## List of abbreviations

GUI: Graphical User Interface
MD: Molecular Dynamics
MeTrEx: Membrane Trajectory Explorer
PMB: Polymyxin B1

## Supplementary information

**File name:** Additional_file_1

- **File format:** .pdf
- **Title of data:** Supplementary Information: MeTrEx: Membrane Trajectory Explorer
- **Description of data:** The screenshots illustrate the features and user interface of the MeTrEx program.

**File name:** Additional_file_2

- **File format:** .xlsx
- **Title of data:** Tool Comparison
- **Description of data:** The full comparison table provides a detailed summary of tools for analysing and visualising MD simulation data.

## Declarations

### Ethics approval and consent to participate

Not applicable.

### Consent for publication

Not applicable.

### Availability of data and materials

The datasets supporting the conclusions of this article are available as open source at Zenodo [29]. MeTrEx, as software, is published on GitHub, and detailed documentation is available to explain a typical workflow and usage. Project name: MeTrEx (Membrane Trajectory Explorer). Project home page: https://github.com/LSI-UniKonstanz/MeTrEx. Operating system(s): Platform independent. Programming language: Python. Other requirements: Python 3.8 or higher and several open-source Python packages: MDAnalysis, MDTraj, NumPy, pandas, scikit-learn, Matplotlib, PyQt6, License: GPL3. Any restrictions to use by non-academics: None.

### Competing interests

The authors declare that they have no competing interests.

### Funding

This work was partly funded by the Deutsche Forschungsgemeinschaft (DFG) under Germany’s Excellence Strategy–EXC 2117–422037984, and DFG project ID 251654672–TRR 161.

### Authors’ contributions

**SJH:** Conceptualisation, Investigation, Writing - Review & Editing. **CR:** Investigation, Visualisation, Software, Writing - Original Draft. **BE:** Investigation, Software, Writing - Original Draft. **KK:** Conceptualisation, Visualisation, Writing - Review & Editing. **YZ:** Conceptualisation, Writing - Review & Editing. **WM:** Conceptualisation, Writing - Review & Editing. **YH:** Investigation, Writing - Review & Editing. **JL:** Conceptualisation, Supervision, Resources, Writing - Review & Editing. **FS:** Conceptualisation, Supervision, Resources, Writing - Review & Editing. All authors read and approved the final manuscript.

## Acknowledgements

Thanks to Martin Kern for providing the code to visualise the membrane as an abstract surface. JL is an Australian National Health and Medical Research Council (NHMRC) Investigator Fellow (APP2025937). WM acknowledges the support provided by the Monash Suzhou Research Institute (MSRI) Scholarship.

## Notes

https://github.com/LSI-UniKonstanz/MeTrEx

https://zenodo.org/records/14512968

### Competing Interest Statement

The authors have declared no competing interest.

