## Additional_file_1 for "MeTrEx: Membrane Trajectory Explorer"

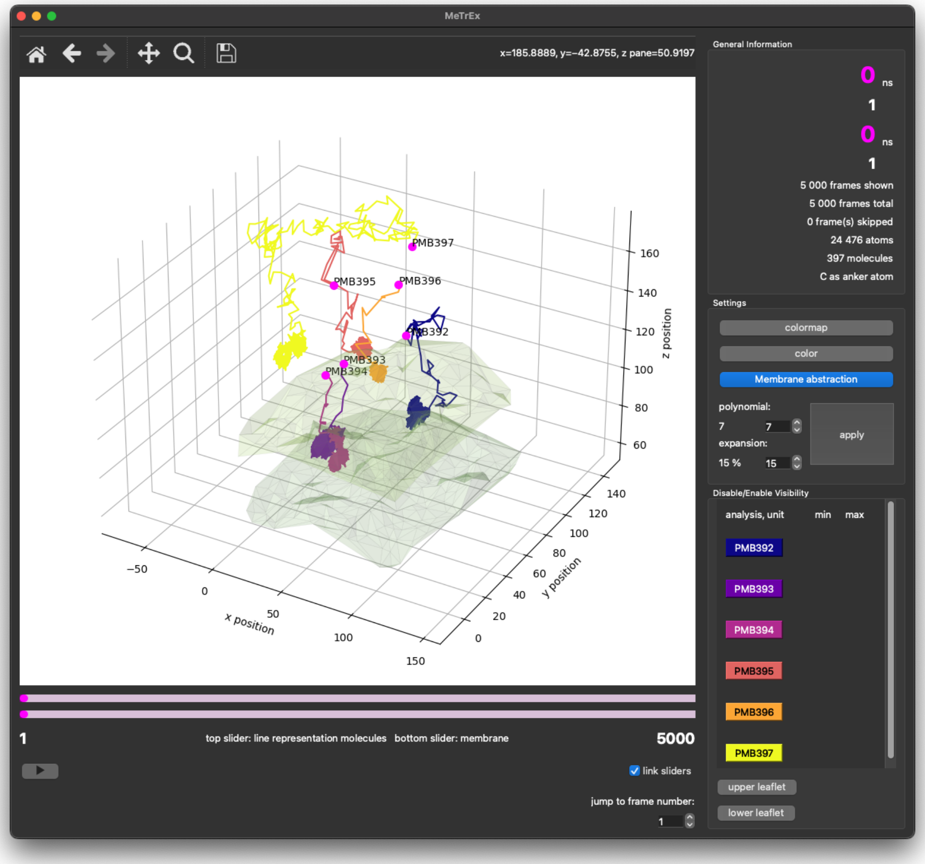


**Figure S1**: The main view displays the trajectories of Polymyxin B1 molecules (PMB), with each molecule labelled by its respective residue number. This visualisation shows molecular and membrane movements over time. The membrane is displayed in its triangulated surface visualisation, using the lipids phosphor atom's positions as vertices. Frames can be selected individually, or a chronological animation of the simulation can be played within the main view. The General Information panel provides precise details, including simulation time in nanoseconds (ns) and other relevant data.

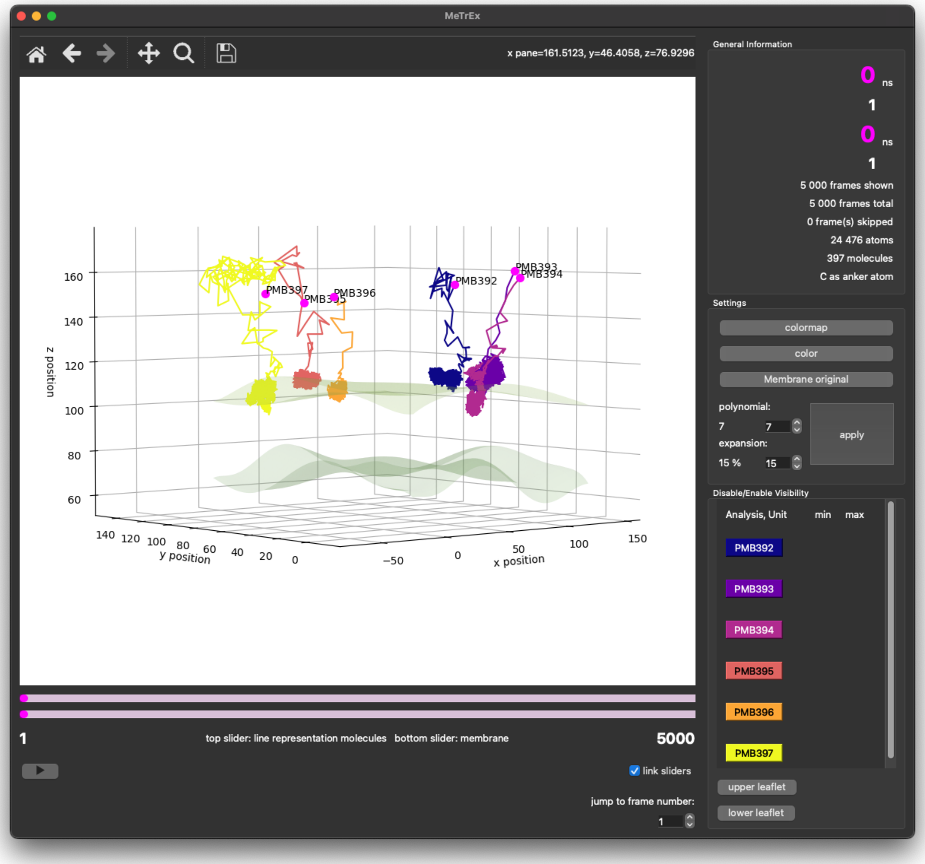


**Figure S2**: MeTrEx allows free movement and rotation of the simulation visualisation, enabling users to observe molecular trajectories from different angles for enhanced analysis of interactions with the membrane surface. Increasing the polynomial degree of the membrane abstraction provides a more detailed and refined representation of the membrane.

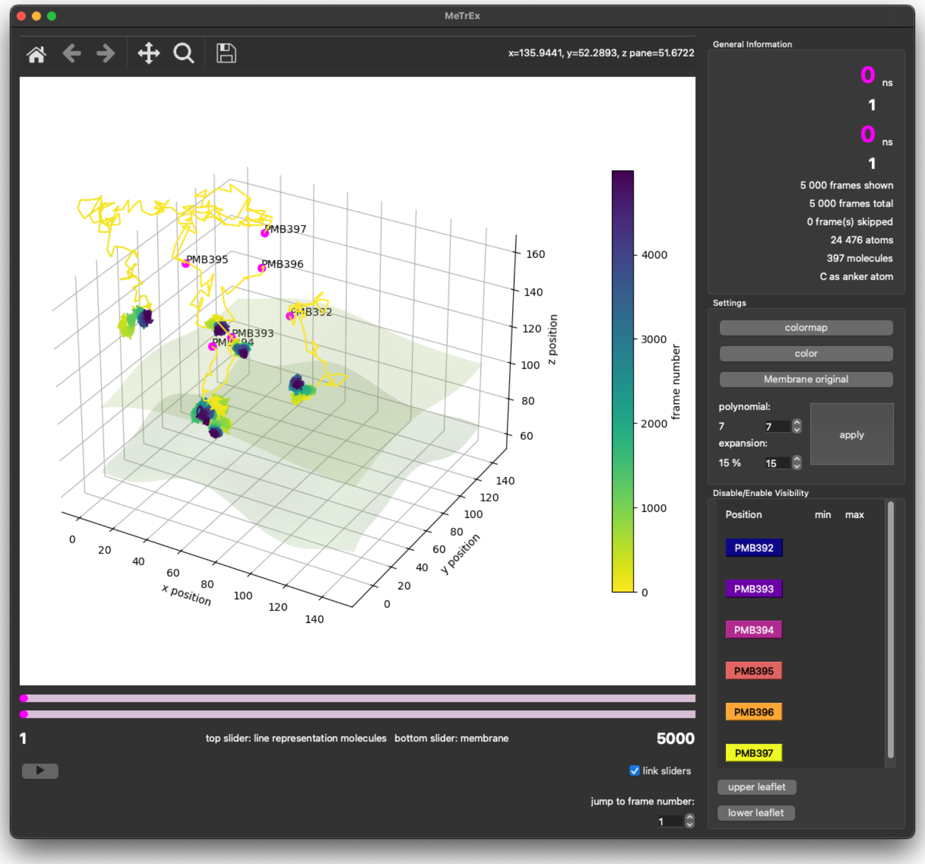


**Figure S3**: The trajectory can be visualised with a colour gradient representing the molecular positions at each frame over time. This allows for a clear depiction of the trajectory's course. Additionally, users can play an animation of the trajectory using the action button for a dynamic, frame-by-frame exploration.

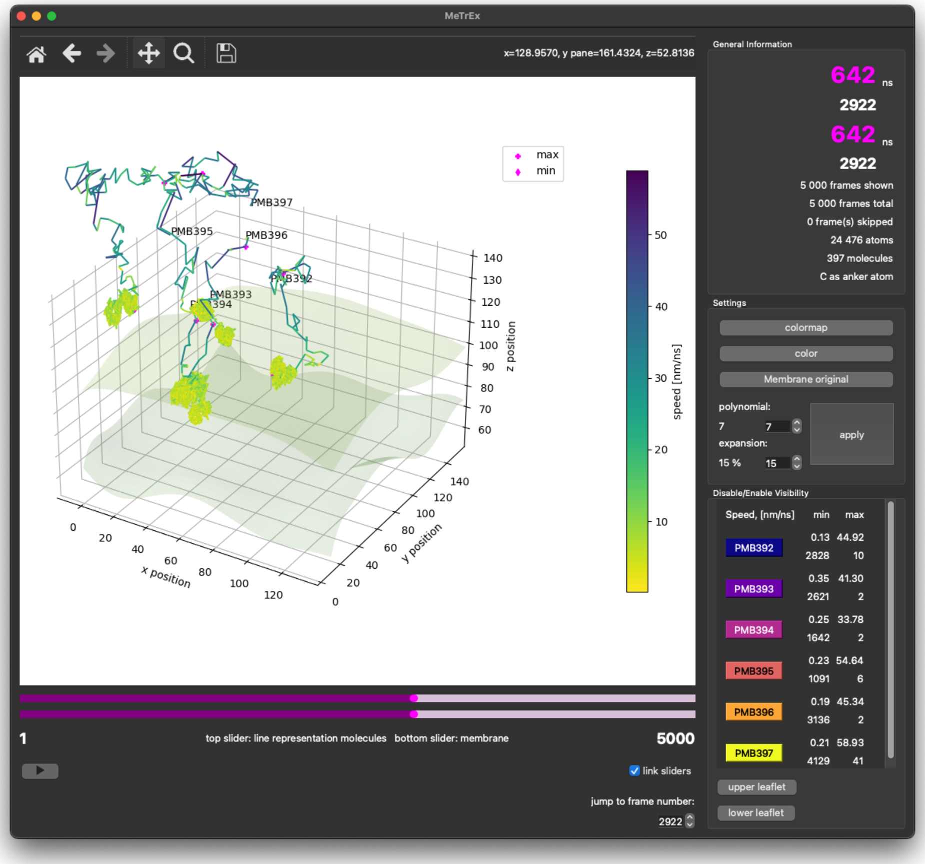


**Figure S4**: Molecular or atomic speed along the trajectories is visualised using a colour gradient, with speeds represented in nm/ns. In the side view, minima and maxima and their corresponding frames are displayed for each analysed molecule. This feature is handy for analysing interactions in ligand-membrane studies, providing insights into molecular dynamics over time.

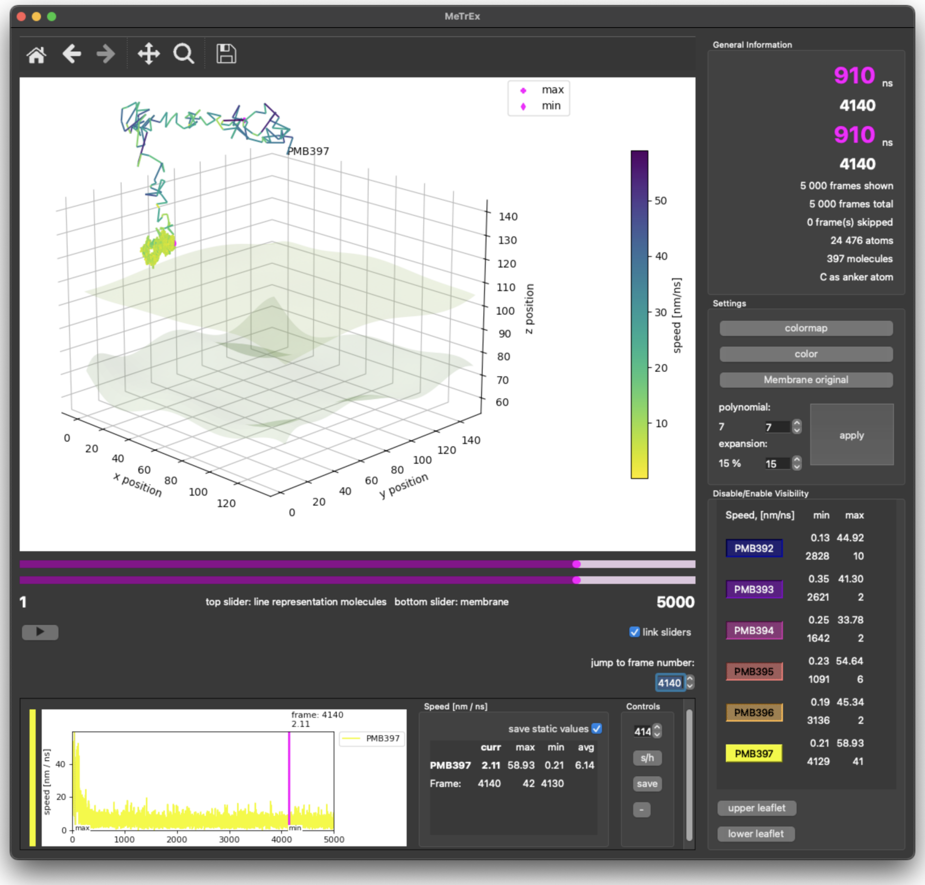


**Figure S5**: Displaying the speed of a single molecule's trajectory in the bottom view: By isolating a selected molecule's trajectory, MeTrEx allows focused analysis. In addition to the colour gradient displayed in the main view, the molecular speed is plotted in the bottom view below, providing detailed information such as speed at the currently selected frame, average speed across the trajectory, and minimum and maximum values.

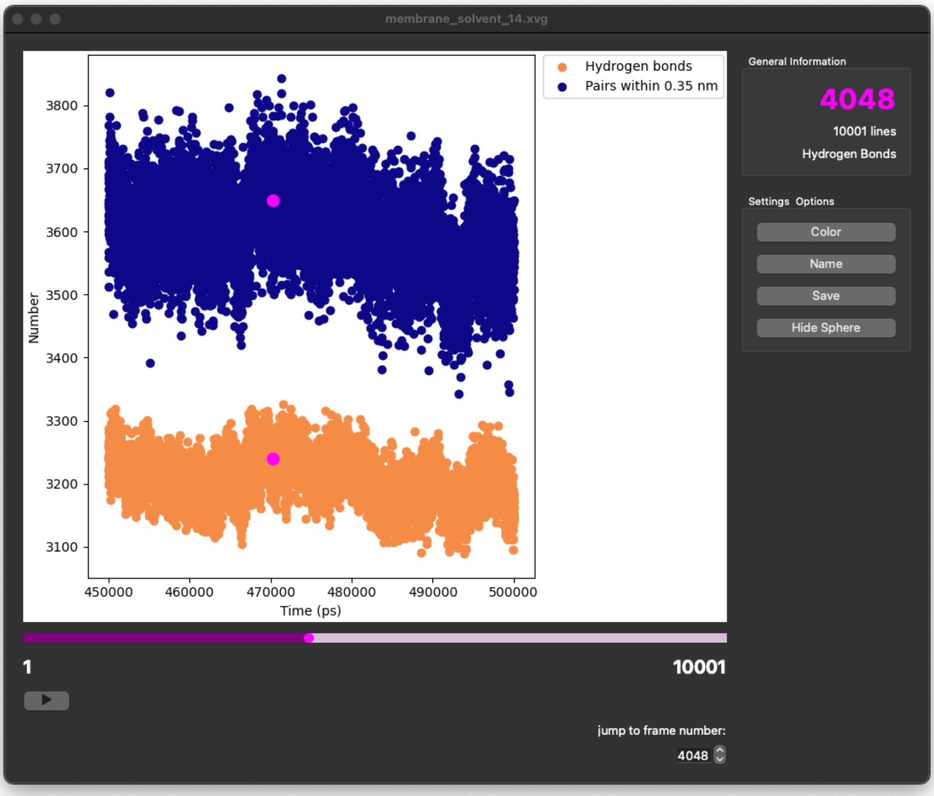


**Figure S6**: The sub-view in MeTrEx offers a streamlined interface for visualising external analysis data in XVG file format. The sub-view shown in the figure features a 2D visualisation, although 3D plots can also be displayed. Each sub-view includes a view area, a slider with related controls beneath it, and an interaction panel on the right. These windows are initiated via the menu option to load external XVG files, with customisable options for colour and data labels. Data from the main view, such as drug molecules, can be linked to the sub-view for enhanced analysis. The sub-view can also be saved as an image file; it provides an info panel displaying the current slider position and total number of frames or lines. All sub-views automatically close when the main window is closed.
